# Dorsoventral comparison of intraspecific polymorphisms in the butterfly wing pattern using a convolutional neural network

**DOI:** 10.1101/2024.08.01.606114

**Authors:** Kai Amino, Tsubasa Hirakasa, Masaya Yago, Takashi Matsuo

**Author notes:** Corresponding author: Kai Amino, Furocho 1, Nagoya, Aichi 464-8603, Japan.

## Abstract

Butterfly wing patterns exhibit notable differences between the dorsal and ventral surfaces, and morphological analyses of them have provided insights into the ecological and behavioural characteristics of wing colour patterns. Conventional methods for dorsoventral comparisons are constrained by the need for homologous patches or shared features between two surfaces, limiting their applicability across species. We used a convolutional neural network (CNN)-based analysis, which can compare images of the dorsal and ventral surfaces without focusing on homologous patches or features, to detect dorsoventral bias in intraspecific polymorphisms such as sexual dimorphism (SD) and female-limited mimetic polymorphism (FMP). Using specimen images of 29 butterfly species from the Yaeyama Islands, Japan, we first showed that the level of SD calculated by CNN-based analysis corresponded well with traditional assessments of SD, demonstrating the validity of the method. Dorsoventral biases were widely detected, particularly in SD, which tended to bias dorsally. This supports the conventional hypothesis that sexual selection acts more strongly on the dorsal surface. In contrast, the FMP analysis showed no significant bias. Our findings highlight CNN-based analysis as a versatile technique for dorsoventral comparisons, suggesting that broader species sampling could reveal general patterns of selection acting differentially on the two surfaces.

## 1. Introduction

Understanding the evolution of the butterfly wing pattern, one of nature’s most diverse colour patterns, has contributed significantly to various fields, including genetics, development, and ecology. One notable feature of the butterfly wings is that the two (dorsal and ventral) surfaces of the wings have distinct colour patterns in many species [1−3]. Such differences are sometimes the focus of research, yielding insights into the ecological and behavioral importance of butterfly wings. For example, Oliver *et al*. [4] compared the evolutionary speed of the appearance of eye spots between the dorsal and ventral surfaces, demonstrating that the number of eye spots evolved most rapidly on the male dorsal surface. Tuomaala *et al*. [5] compared melanization between the two surfaces and found that melanization along latitudes was not seen only on the male dorsal surface. Based on their findings, they concluded that the sexually different selections (i.e., sexual selection) acts more strongly on the dorsal surface than on the ventral surface. However, because their research was restricted to the genus level at most [4], such analyses across a broader range of species are needed to strengthen the credibility of their conclusions.

Conventional analyses that compare dorsal and ventral wing patterns (“dorsoventral comparisons”) can be divided into two types. One method is patch comparison, which compares the shapes or colours of patches that are homologous between the dorsal and ventral surfaces (for example, patches of the same colours, shapes, or positions on both surfaces). Tuomaala *et al*. [5] dorsoventrally compared the spectral pattern of white base colour in *Pieris napi*, whereas Klein and Araujo [6] compared the size of red colour patches in *Heliconius* butterflies. The second approach is feature comparison, which dorsoventrally compares the properties of certain features shared by both surfaces. Oliver et al. [4] focused on the number of eye spots, and Rutowski et al. [7] focused on iridescence (the change in colour reflectance with direction). However, each of these approaches has limitations when applied to a wider range of species. Patch comparisons are not possible when homologous patches do not exist (for example, in some nymphalid butterflies with dorsoventrally different colour components [1, 2]). On the other hand, feature comparison is limited to species that share certain features, such as eye spots. These constraints may have precluded dorsoventral comparisons from being used among a wider range of species, such as across families or genera.

The development of deep learning using convolutional neural networks (CNNs) has revolutionized computer vision technology. They can perform a variety of supervised learning tasks, including image classification and object detection, which can also be used for biological image analysis [8–10]. Furthermore, CNNs trained for such supervised tasks have been shown to have sophisticated feature extraction capabilities. For example, a technique known as “transfer learning” allows CNNs to perform well even when they are not trained on the objects they are classifying. Similarly, the intermediate output of a CNN pre-trained on a large-scale dataset has been shown to be useful in assessing the similarity between input images, even when CNNs are not trained for those input images [11, 12]. Pre-trained CNNs have been shown to measure similarities successfully on biological images, such as bumble bees [13] and fish body texture [14]. The CNN-based similarity measurement appear to solve the difficulty in dorsoventral comparison of butterfly wing patterns. It can calculate the similarity between colour patterns by simply inputting images, and it does not need to focus on certain colour patches or features that are homologous between two surfaces. As a result, CNN is considered to be a credible method for studying the dorsoventral patterns of butterfly wings.

Here we tested the usability of a CNN-based analysis for the dorsoventral comparison in butterfly wing patterns by determining whether dorsoventral bias can be detected in two types of intraspecific polymorphisms: sexual dimorphism (SD) and female-limited mimetic polymorphism (FMP). SD, in which males and females exhibit distinct morphological differences, is commonly observed among butterfly wing patterns [15] and is considered the result of sexual selection [16], such as mate choice. Previous research has suggested that mate choice acts on the dorsal surface, which is exposed during butterfly courtship displays [17, 18], perhaps resulting in a higher level of SD (i.e., dissimilarity between sexes) on the dorsal surface. The latter FMP, in which only a portion of females of nontoxic species resemble other toxic species but males and other females do not, can be found in papilionid butterflies [19–21]. Because predation pressure, the putative primary cause of FMP evolution [22], is thought to act on the dorsal surface, which is visible to aerial predators (i.e., birds) when butterflies are flying [23], we expected to detect a dorsal bias in the level of FMP.

## 2. Materials and methods

### 2.1. Image acquisition and pre-processing

We used a total of 29 butterfly species observed in the Yaeyama Islands, Japan [24]. These butterflies cover 20 genera from two families, Nymphalidae and Papilionidae (table S1), and include species with varying levels of SD on both wing surfaces for several traits such as base colour, spot colour, eye spots, and stripes (figure 1*a*), as well as two species with FMP (figure 1*b*; [25, 26]). Furthermore, SD in *Hypolimnas misippus, Argyreus hyperbius*, and *Euploea mulciber* may be linked to Batesian or Müllerian mimicry (table S1; [27–30]), providing a better understanding of the interactive effect of sexual selection and predation pressure on dorsoventral bias in SD.

**Figure 1.**
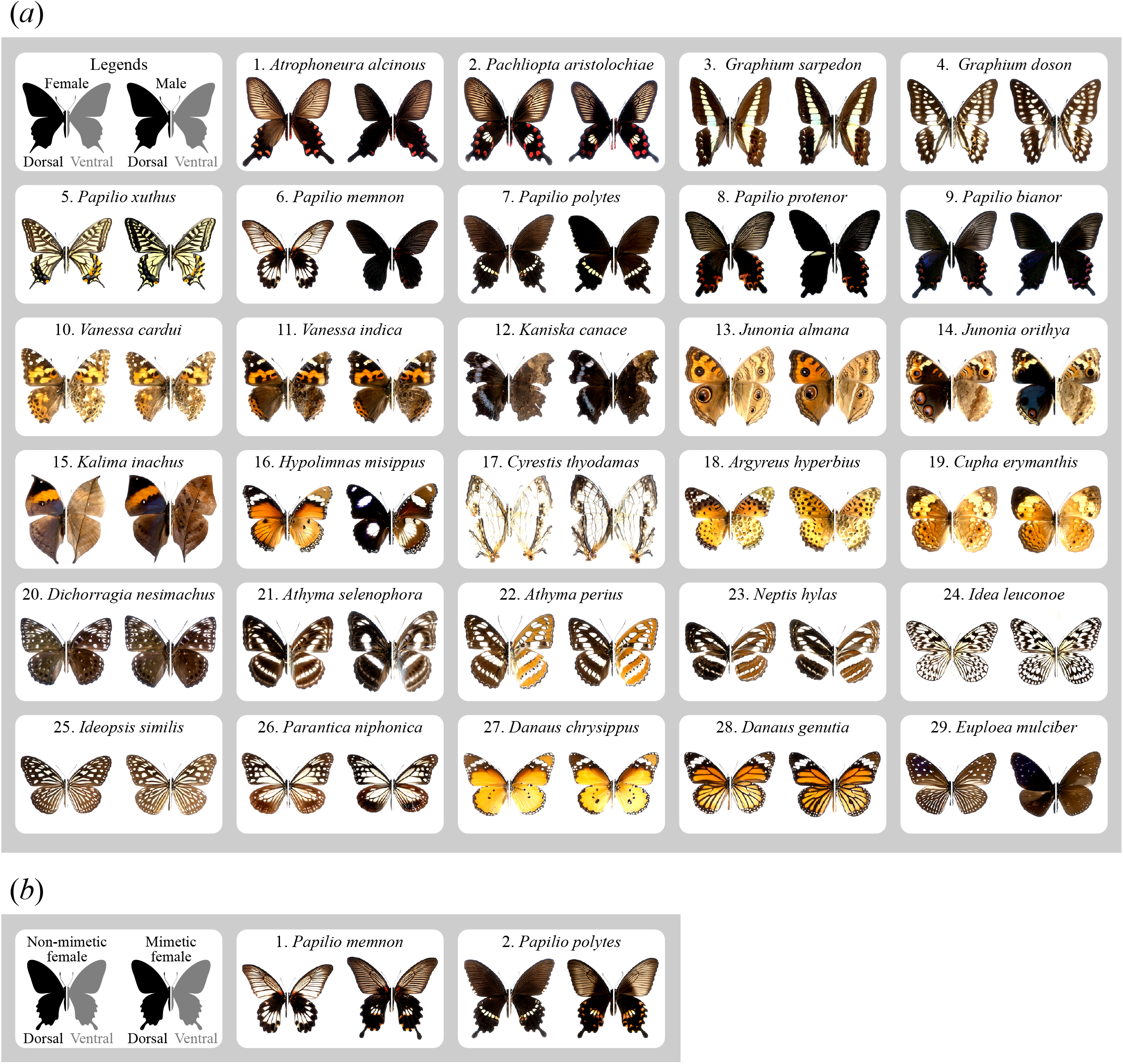
Images of dorsal and ventral wing surfaces of the butterfly species used for (*a*) SD and (*b*) FMP analyses.

We photographed these butterflies from a large butterfly specimen dataset “Suguru Igarashi Insect Collection” (e.g., [31]), stored in the University Museum, The University of Tokyo (UMUT). To minimise individual variation, 3-5 individuals per sex or morph (mimetic or non-mimetic) were photographed for each species (table S1), yielding a total of 592 images for the entire image dataset (table S2). Furthermore, for each species, samples included specimens collected in as close proximity as possible to prevent confounding the effects of intraspecific variation with sampled sites. For this purpose, specimens collected outside of the Yaeyama Islands were also used for some species.

When photographing the specimens, they were placed on a white background and illuminated from the side to prevent the shadow of the specimen from casting on the background, ensuring uniformity of background information (figure S1*a*). A digital camera (E-M5MarkIII, Olympus, Tokyo, Japan) with a zoom lens (M.ZUIKO DIGITAL ED 40-150mm F2.8; Olympus) was mounted approximately 50 cm above the specimen to capture the images. Original RAW photos (ORF, Olympus RAW File) were exported as PNG files. The following preprocessing steps were performed using Adobe Photoshop. To reduce differences in light conditions, colour correction was performed based on the black and white points of the colour chart, which were captured in each specimen photograph (figure S1*b*). Images were trimmed to fit the individual in the centre of a square frame of 224*224 px (the standard input size for CNN [32]), reducing the effect of body size differences among individuals (figure S1*c*). The photographic dataset used for our analysis is provided in the Dryad Data Repository with filenames corresponding to joined data in table S2.

### 2.2. Assessing wing pattern distance using CNN-based analysis

We used the previously developed metric for similarity measurement called “Learned Perceptual Image Patch Similarity (LPIPS)”, following the methods described by Zhang et al. [33]. When two images are passed through a CNN (AlexNet; [34]) that has already been trained on a large-scale image recognition dataset (ImageNet; [32]), the activations from the convolutional layers, which are numerical (more than 1000) representations of the images, are calculated. LPIPS is then computed by linearly rescaling the Euclidian distance between these activation vectors of two images, yielding “zero” for identical images and a higher score for a pair of images with lower similarity [33]. Although LPIPS is calculated by a CNN trained on ImageNet [32], which contains a variety of objects other than organisms, such as food, people, and vehicles, it has also been shown to reflect the similarity between images of a wide range of organisms [35] and produce more accurate similarity than the conventional approach [13, 35, 36].

### 2.3. Statistical analysis

To confirm the validity of the CNN-based similarity measurement, we first compared the level of SD in wing patterns calculated by the CNN (dorsal + ventral) with the description in a pictorial book [37] about the difficulty in distinguishing sexes (table 1), which should be the ground truth for the similarity between sexes for each species. In addition, we defined the rank of SD (RSD) from 1 to 4 based on the description (table 1), enabling the quantitative evaluation of the concordance between the ground truth and the SD calculated by the CNN. The Steel-Dwass test was used to determine the statistical significance of the difference in SD levels between the groups of different RSD. We also used an ordered logistic regression to test whether the RSD was positively correlated with that calculated by the CNN.

**Table 1.**
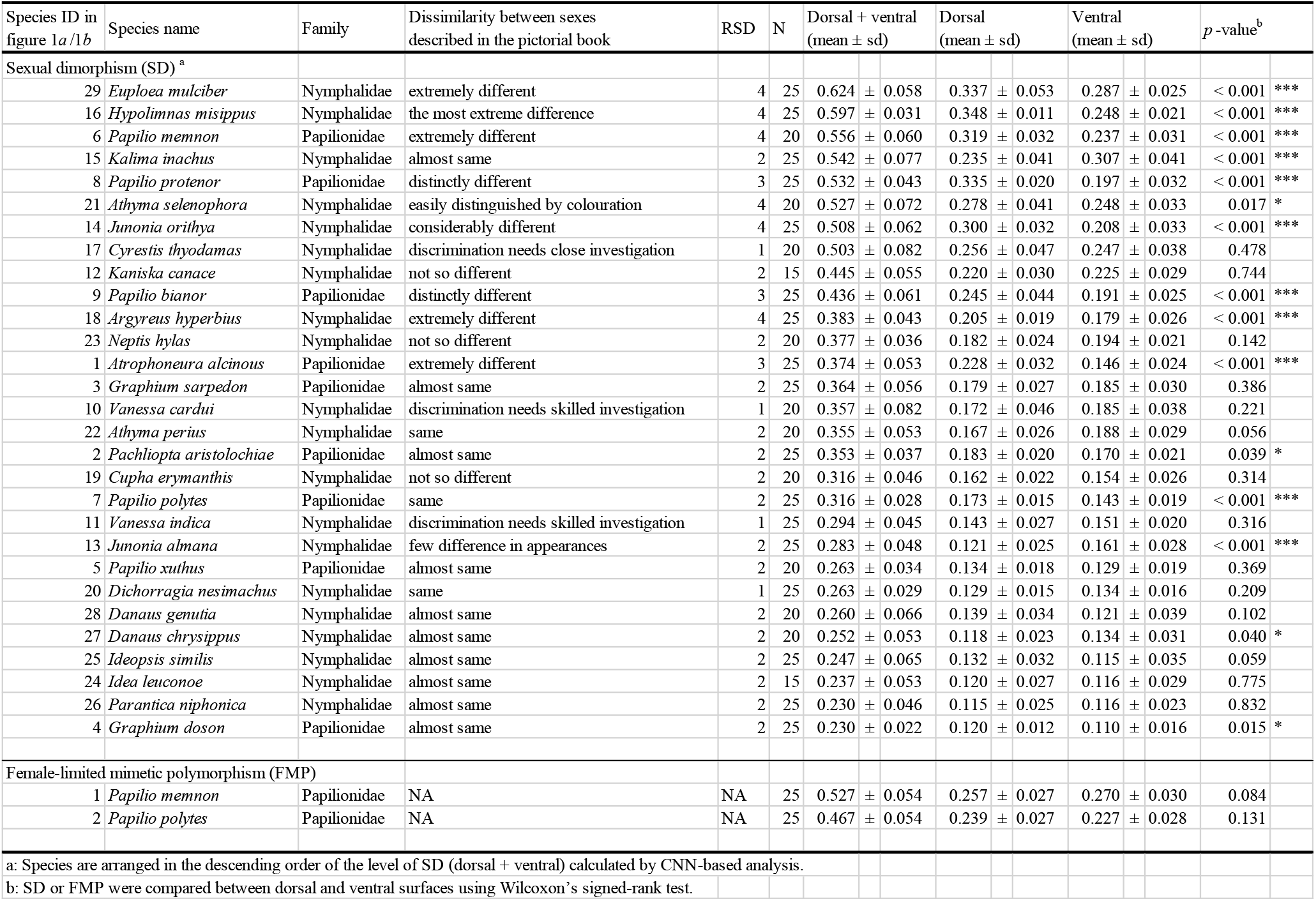
Dorsoventral comparisons of intra-specific polymorphisms for each species.

|  |  |  |  |  |  |  |  |  |  |  |
| --- | --- | --- | --- | --- | --- | --- | --- | --- | --- | --- |
| Species ID in figure 1a/1b | Species name | Family | Dissimilarity between sexes described in the pictorial book | RSD | N | Dorsal + ventral (mean ± sd) | Dorsal (mean ± sd) | Ventral (mean ± sd) | *p*-value<sup>b</sup> | |
| Sexual dimorphism (SD)<sup>a</sup> |  |  |  |  |  |  |  |  |  |  |
| 29 | *Euploea mulciber* | Nymphalidae | extremely different | 4 | 25 | 0.624 ± 0.058 | 0.337 ± 0.053 | 0.287 ± 0.025 | < 0.001 | *** |
| 16 | *Hypolimnys misippus* | Nymphalidae | the most extreme difference | 4 | 25 | 0.597 ± 0.031 | 0.348 ± 0.011 | 0.248 ± 0.021 | < 0.001 | *** |
| 6 | *Papilio memnon* | Papilionidae | extremely different | 4 | 20 | 0.556 ± 0.060 | 0.319 ± 0.032 | 0.237 ± 0.031 | < 0.001 | *** |
| 15 | *Kalima inachus* | Nymphalidae | almost same | 2 | 25 | 0.542 ± 0.077 | 0.235 ± 0.041 | 0.307 ± 0.041 | < 0.001 | *** |
| 8 | *Papilio protenor* | Papilionidae | distinctly different | 3 | 25 | 0.532 ± 0.043 | 0.335 ± 0.020 | 0.197 ± 0.032 | < 0.001 | *** |
| 21 | *Athyma selenophora* | Nymphalidae | easily distinguished by colouration | 4 | 20 | 0.527 ± 0.072 | 0.278 ± 0.041 | 0.248 ± 0.033 | 0.017 | * |
| 14 | *Junonia orithya* | Nymphalidae | considerably different | 4 | 25 | 0.508 ± 0.062 | 0.300 ± 0.032 | 0.208 ± 0.033 | < 0.001 | *** |
| 17 | *Cyrestis thyodamas* | Nymphalidae | discrimination needs close investigation | 1 | 20 | 0.503 ± 0.082 | 0.256 ± 0.047 | 0.247 ± 0.038 | 0.478 | |
| 12 | *Kaniska canace* | Nymphalidae | not so different | 2 | 15 | 0.445 ± 0.055 | 0.220 ± 0.030 | 0.225 ± 0.029 | 0.744 | |
| 9 | *Papilio bianor* | Papilionidae | distinctly different | 3 | 25 | 0.436 ± 0.061 | 0.245 ± 0.044 | 0.191 ± 0.025 | < 0.001 | *** |
| 18 | *Argyreus hyperbius* | Nymphalidae | extremely different | 4 | 25 | 0.383 ± 0.043 | 0.205 ± 0.019 | 0.179 ± 0.026 | < 0.001 | *** |
| 23 | *Neptis hylas* | Nymphalidae | not so different | 2 | 20 | 0.377 ± 0.036 | 0.182 ± 0.024 | 0.194 ± 0.021 | 0.142 | |
| 1 | *Atrophaneura alcinous* | Papilionidae | extremely different | 3 | 25 | 0.374 ± 0.053 | 0.228 ± 0.032 | 0.146 ± 0.024 | < 0.001 | *** |
| 3 | *Graphium sarpedon* | Papilionidae | almost same | 2 | 25 | 0.364 ± 0.056 | 0.179 ± 0.027 | 0.185 ± 0.030 | 0.386 | |
| 10 | *Vanessa cardui* | Nymphalidae | discrimination needs skilled investigation | 1 | 20 | 0.357 ± 0.082 | 0.172 ± 0.046 | 0.185 ± 0.038 | 0.221 | |
| 22 | *Athyma perius* | Nymphalidae | same | 2 | 20 | 0.355 ± 0.053 | 0.167 ± 0.026 | 0.188 ± 0.029 | 0.056 | |
| 2 | *Pachliopta aristolochiae* | Papilionidae | almost same | 2 | 25 | 0.353 ± 0.037 | 0.183 ± 0.020 | 0.170 ± 0.021 | 0.039 | * |
| 19 | *Cupha erymanthis* | Nymphalidae | not so different | 2 | 20 | 0.316 ± 0.046 | 0.162 ± 0.022 | 0.154 ± 0.026 | 0.314 | |
| 7 | *Papilio polytes* | Papilionidae | same | 2 | 25 | 0.316 ± 0.028 | 0.173 ± 0.015 | 0.143 ± 0.019 | < 0.001 | *** |
| 11 | *Vanessa indica* | Nymphalidae | discrimination needs skilled investigation | 1 | 25 | 0.294 ± 0.045 | 0.143 ± 0.027 | 0.151 ± 0.020 | 0.316 | |
| 13 | *Junonia almana* | Nymphalidae | few difference in appearances | 2 | 25 | 0.283 ± 0.048 | 0.121 ± 0.025 | 0.161 ± 0.028 | < 0.001 | *** |
| 5 | *Papilio xuthus* | Papilionidae | almost same | 2 | 20 | 0.263 ± 0.034 | 0.134 ± 0.018 | 0.129 ± 0.019 | 0.369 | |
| 20 | *Dichorragia nesimachus* | Nymphalidae | same | 1 | 25 | 0.263 ± 0.029 | 0.129 ± 0.015 | 0.134 ± 0.016 | 0.209 | |
| 28 | *Danaus genutia* | Nymphalidae | almost same | 2 | 20 | 0.260 ± 0.066 | 0.139 ± 0.034 | 0.121 ± 0.039 | 0.102 | |
| 27 | *Danaus chrysippus* | Nymphalidae | almost same | 2 | 20 | 0.252 ± 0.053 | 0.118 ± 0.023 | 0.134 ± 0.031 | 0.040 | * |
| 25 | *Ideopsis similis* | Nymphalidae | almost same | 2 | 25 | 0.247 ± 0.065 | 0.132 ± 0.032 | 0.115 ± 0.035 | 0.059 | |
| 24 | *Idea leuconoe* | Nymphalidae | almost same | 2 | 15 | 0.237 ± 0.053 | 0.120 ± 0.027 | 0.116 ± 0.029 | 0.775 | |
| 26 | *Parantica niphonica* | Nymphalidae | almost same | 2 | 25 | 0.230 ± 0.046 | 0.115 ± 0.025 | 0.116 ± 0.023 | 0.832 | |
| 4 | *Graphium doson* | Papilionidae | almost same | 2 | 25 | 0.230 ± 0.022 | 0.120 ± 0.012 | 0.110 ± 0.016 | 0.015 | * |
| Female-limited mimetic polymorphism (FMP) |  |  |  |  |  |  |  |  |  |  |
| 1 | *Papilio memnon* | Papilionidae | NA | NA | 25 | 0.527 ± 0.054 | 0.257 ± 0.027 | 0.270 ± 0.030 | 0.084 | |
| 2 | *Papilio polytes* | Papilionidae | NA | NA | 25 | 0.467 ± 0.054 | 0.239 ± 0.027 | 0.227 ± 0.028 | 0.131 | |
| a: Species are arranged in the descending order of the level of SD (dorsal + ventral) calculated by CNN-based analysis. |  |  |  |  |  |  |  |  |  |  |
| b: SD or FMP were compared between dorsal and ventral surfaces using Wilcoxon's signed-rank test. |  |  |  |  |  |  |  |  |  |  |

Next, we compared the levels of intraspecific polymorphisms between dorsal and ventral surfaces. The SD and MP were calculated for all possible pairs in each species (for example, 25 pairs of SD if five males and five females were available). To test whether the CNN-based analysis could detect dorsoventral bias in intraspecific polymorphisms, we used Wilcoxon’s signed rank test to compare the SD and FMP between the dorsal and ventral surfaces of each species. The tendency for dorsoventral bias was then examined in groups divided according to the types of selection (sexual selection or predation pressure) putatively involved in the evolution of the polymorphism. For SD, we first divided the species based on whether the SD was linked to mimicry (table S1), to avoid confusing the effect of sexual selection with predation pressure. Species in which SD was not linked to mimicry were also classified according to RSD (1 – 4), with the expectation that the dorsoventral bias of SD would increase with the level of SD if our analysis was correct. We then used the Wilcoxon’s signed-rank test to assess the level of SD in five groups (one mimicry-related and four mimicry-non-related groups) and the level of FMP between the dorsal and ventral surfaces.

## 3. Results and discussion

### 3.1. The accuracy of CNN-based analysis

Table 1 shows the species arranged in descending order of the level of SD calculated by the CNN, referring to the difficulties in distinguishing between the sexes described in the pictorial book. The rank of the species where the sexes are easy to distinguish (described as “apparently different” or “extremely different”) tends to be high (table 1: *E. mulciber, H. misippus, Papilio memnon*), whereas the rank of the species where the sexes are difficult to distinguish (described as “almost same”) tends to be low (table 1: *Graphium doson, Parantica niphonica*). Moreover, the level of SD calculated by CNN was significantly different between the species with different RSD (figure 2*a*), and the correlation between them was significantly positive (*p* < 0.001, coefficient = 10.71 by ordered logistic regression). These results suggest that the CNN accurately calculated the SD levels of the butterfly specimen images.

**Figure 2.**
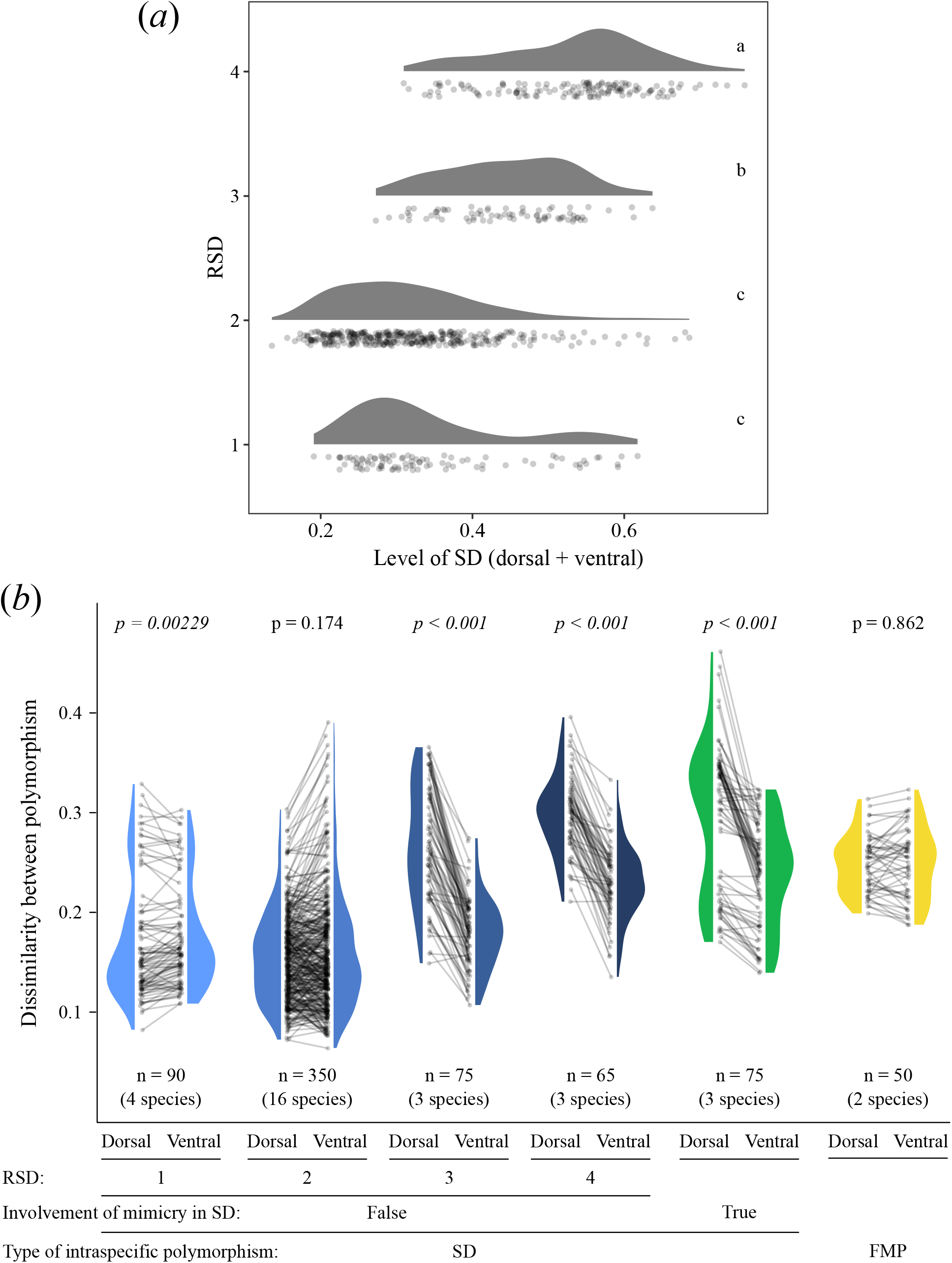
(*a*) The relationship between the level of SD calculated by CNN-based analysis and RSD. Violin plots with the same characters are not significantly different by the Steel-Dwass test. (*b*) Dorsoventral comparisons of SD and FMP. The pair of grey dots connected by the line represents the dorsal and ventral SD (or FMP) of each pair. The *p*-values were calculated by Wilcoxon’s signed-rank test.

In previous studies providing the validity of similarity measurements by CNN, the use of the ground - truth similarity data, such as similarity judgements made by human volunteers, was preferred [11, 12, 33]. However, the validity of similarity assessments in biological studies, where ground-truth data are seldom available, relies solely on the accuracy of species or subspecies classification [9, 14], which may not always reflect similarity. We solved this problem by using the pictorial book’s description of the difficulties of discriminating sexes as ground-truth data, which provided stronger evidence for the accuracy of similarity measurements using CNN.

### 3.2. Dorsoventral bias in intraspecific polymorphism

Dorsoventral biases in intraspecific polymorphisms were significantly detected in many species (table 1). Dorsoventral bias in SD was detected even in papilionid butterflies, where the difference between the dorsal and ventral wing surfaces was moderate (figure 1*a*; 1-9), demonstrating the sensitivity of our method. Moreover, *Junonia orithya* (figure 1*a*; 14) showed dorsoventral bias in SD, indicating that our method can be broadly applied to such species with completely different colour pattern components between two surfaces. In this species, conventional colour comparison between dorsoventrally homologous patches has been difficult to apply (blue structural base colour and large eye spots on dorsal hindwings, with tree-bark mimesis pattern and small eye spots on ventral hindwings).

For SD not related to mimicry (figure 2*b*, blue), we found no significant tendency for bias or a significant but small bias on the ventral surface in groups exhibiting a lower level of SD (RSD = 2, 1, respectively), whereas in groups with a higher level of SD (RSD = 3, 4), we found that SD was biased on the dorsal surface. This suggests that SD tends to bias on the dorsal surface, which supports the conventional view that the dorsal surface, which is utilized for sexual display, is the primary site of sexual selection [17, 18]. Our results indicate that the dorsal bias in sexual selection previously identified in the genus *Bicyclus* [4] and the species *P. napi* [5] can be found in a wide range of species from two families, Papilionidae and Nymphalidae. Although we expected dorsal bias in FMP based on the previous hypothesis suggesting the dorsally biased predation pressure on mimetic wing patterns [23], no significant difference was seen (figure 2*b*, yellow). Because the number of species available in Japan for FMP study was limited, this result does not necessarily imply that there was no bias in the level of FMP on either surface. The SD related to mimicry, which can be attributed to both sexual selection and predation pressure, was significantly biased toward the dorsal surface (figure 2*b*, green). This result appears acceptable by combining the foregoing results suggesting that SD is biased on the dorsal surface and FMP is scarcely biased on either surface.

### 3.3. Future directions

Our study aimed to examine the versatility of CNN-based analysis in the dorsoventral comparison of butterfly wing patterns and found that our approach can be applied to species for which conventional methods are difficult to apply. For this purpose, we chose 29 species living in the Yaeyama Islands as the minimal number of samples, which cover a wider range of species than prior research and vary in the degree of dorsoventral and intraspecific differences in wing patterns. Although the dorsoventral bias in SD and FMP may provide insights into the selections that contribute to the evolution of such intraspecific polymorphisms, we need to increase the number and diversity of species to determine the general tendency of dorsoventral bias in selective pressures.

Although our study demonstrates the usefulness of CNN-based analysis for investigating dorsoventral disparities in butterfly wing patterns, caution is required when interpreting the results as the evidence of dorsoventral bias in selective pressure alone. For example, developmental coupling between dorsal and ventral wing pattern features (e.g., colour patch size [6] and iridescent colour [7]) might result in dorsoventrally equal expression of a trait which is favoured on one surface but not on the other. In our study, for example, there was no significant difference in FMP levels between two surfaces (figure 2b), which might be interpreted as developmental coupling, as well as the dorsoventrally comparable level of predation pressure. Previous research has investigated inter-individual correlation [38] and response to artificial selection [39] to quantify the strength of the developmental coupling between wing pattern features. These methods, as well as a dorsoventral comparison of wing colour patterns, are needed to understand dorsoventral differences in selective pressure on butterfly wings.

