## Supplementary figure 1 for "Dorsoventral comparison of intraspecific polymorphisms in the butterfly wing pattern using a convolutional neural network"

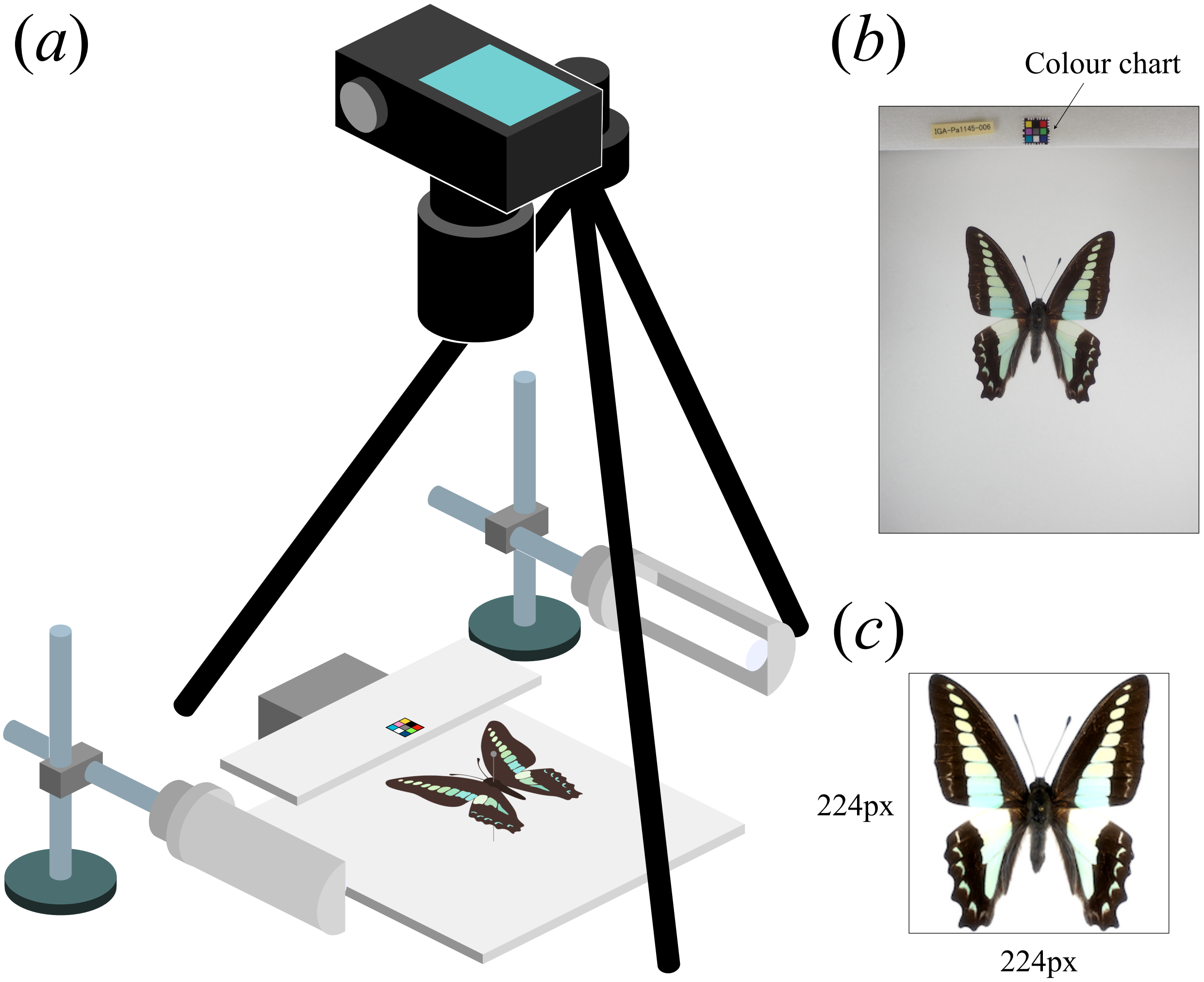


Supplementary figure 1. (*a*) The set up for taking images of butterfly specimens. The colour chart was set at the same height as the wing surface of the specimen. (*b*) The raw image. (*c*) The image for input pre-processed through retouching, tilt correction, and cropping.
